# Plasma Membrane Damage by Environmental Materials Enhances Cell-Cell Fusion and Impairs Immune Functions of Macrophages

**DOI:** 10.1101/2025.06.17.660197

**Authors:** Dandan Wu, Xing Zhang, Ruoning Wang, Amanda Livingston, Eliane El Hayek, Xiang Xue, Aaron Neumann, Matthew Campen, Meilian Liu, Xuexian O. Yang

## Abstract

Macrophages are the most abundant phagocytes and play an essential role in host defense. Previous studies have shown that many environmental materials can activate macrophages and trigger inflammatory responses. However, whether these exposures alter macrophage function in host defense remains unclear. This study found that many environmental materials (such as carbon nanotubes, tungsten carbide [WC], and detergents) can damage the plasma membranes of macrophages. This damage leads to decreased reactive oxygen species (ROS) production and phagocytosis but elevated cell-cell fusion. In vivo, airway exposure to laundry detergent impaired the recruitment of macrophages and other myeloid cells to the lung and dramatically dampened protective TH1 and TH17 cell responses, leading to increased susceptibility to Candida infection in mice. Overall, our data indicate that exposure to environmental materials compromises macrophage membrane integrity and impairs host defense. These findings may aid in the development of effective preventive and therapeutic strategies.

## Introduction

Both man-made and natural materials have been widely used since the Industrial Revolution. Many of these materials are harmful and pose risks to human health. Among environmental hazards, particulate matter (PM) is particular concern due to its surface properties, including size, shape, and chemical composition. Air pollutants contain PM of various-sizes, with PM2.5 and PM0.1 being especially harmful, as they can be inhaled deeply into the lower respiratory tract, traverse the epithelial barrier, and reach multiple organs. PM exposure significantly impacts human health, contributing to the development of respiratory, cardiovascular, and neurological diseases, as well as increasing the risk of certain cancers ^1^.

Tungsten (in its carbide form, or WC) and cobalt are hard metals widely used in cutting and drilling tools, abrasives, and wear-resistant coatings. Occupational exposure to hard metal dusts, such as WC and cobalt, has been associated with respiratory disorders, skin allergies, and even lung cancer in industrial workers ^2–5^. Carbon nanotubes—cylindrical nanostructures composed of carbon atoms—are valued for their exceptional mechanical, electrical, and thermal properties, making them attractive for applications in electronics, materials science, and medicine ^6^. However, certain types of carbon nanotubes, when inhaled, can induce inflammation, oxidative stress, and lung damage ^7,8^. In addition to PM, soluble substances, such as surfactants and phosphates (components of common detergents), can be toxic upon ingestion or skin absorption ^9–12^. Inhalation of detergent residues can cause bronchoconstriction and airway hyperresponsiveness, increasing the risk of asthma attacks. Chronic exposure to these chemicals may lead to adverse long-term health effects, including organ damage.

Macrophages play a crucial role in the immune defense and tissue homeostasis, especially within the innate immune system. They are responsible for detecting, engulfing, and degrading pathogens, foreign particles, and cellular debris ^13^. Environmental materials, such as tungsten carbide, carbon nanotubes, and certain detergent components have been shown to alter macrophage function, including phagocytosis, inflammatory response, and cytotoxicity ^14–16^, potentially compromising host defense mechanisms. Macrophages not only act as individual cells but also form multinucleated giant cells (MGCs), which are functionally distinct from conventional macrophages ^17–19^. Based on their anatomical location, morphology, and function, MGCs can be classified into osteoclasts (specialized in bone remodeling), Langhans giant cells (LGCs; involved in of infectious and non-infectious granulomatous diseases), and foreign body giant cells (FBGCs; present at the site of prosthetic implantation or medical device placement). Notably, occupational or environmental exposure to hard metals can cause hard metal lung disease, also known as giant cell interstitial pneumonia ^20–24^. Despite these associations, the mechanisms by which environmental hazards alter macrophage function and contribute to disease are not fully understood.

A better understanding of the interactions between macrophages and environmental materials is critical for evaluating health risks and developing effective interventions. In this study, demonstrate that various environmental substances—including laundry detergent, saponin (a natural surfactant), carbon nanotubes, WC, and kaolin—can cause plasma membrane damage in macrophages. This damage promotes MGC formation through a CD44-dependent mechanism. Moreover, detergent and saponin treatment suppress reactive oxygen species (ROS) production and impair macrophage phagocytic activity. In vivo, airway exposure to detergent impairs antifungal immune responses in mice. Together, our findings suggest that environmental materials compromise macrophage membrane integrity and protective immune function. These insights may help us develop novel preventative and therapeutic options.

## Results

### Environmental materials damage the plasma membrane of macrophages

Macrophages are the most abundant phagocytes residing in the skin, airway, and gut, playing a crucial role in host defense and tissue homeostasis. As foreign materials, environmental matters can activate macrophages and initiate inflammatory responses. However, it is unclear whether such exposures affect macrophage function and impair host defense. To investigate the effects of environmental materials on macrophages, we first examined the integrity of macrophage cell membranes following exposure to several environmental substances. Briefly, bone marrow-derived macrophages (BMDMs) were treated with various environmental agents or a vehicle control. The cells were then stained with Annexin V (binding to intramembrane-specific lipids to detect membrane damage) and propidium iodide (PI, staining DNA in apoptotic/necrotic cells). Detergents are broadly used to lyse cells in biological research. As expected, treatment with detergent or another surfactant saponin damaged the membranes of macrophages compared to the vehicle treatment **(Figure 1A)**. Treatments with nanoparticles (carbon nanotubes, WC, and kaolin) also disrupted macrophage membranes **(Figure 1B**, **Supplemental Figure 1)**. Additionally, the treatments slightly increased cell death, as revealed by PI staining **(Figure 1**, **Supplemental Figure 1)**. These findings suggest that the tested environmental materials can compromise macrophage membranes, and therefore, may influence macrophage function.

**Figure 1.**
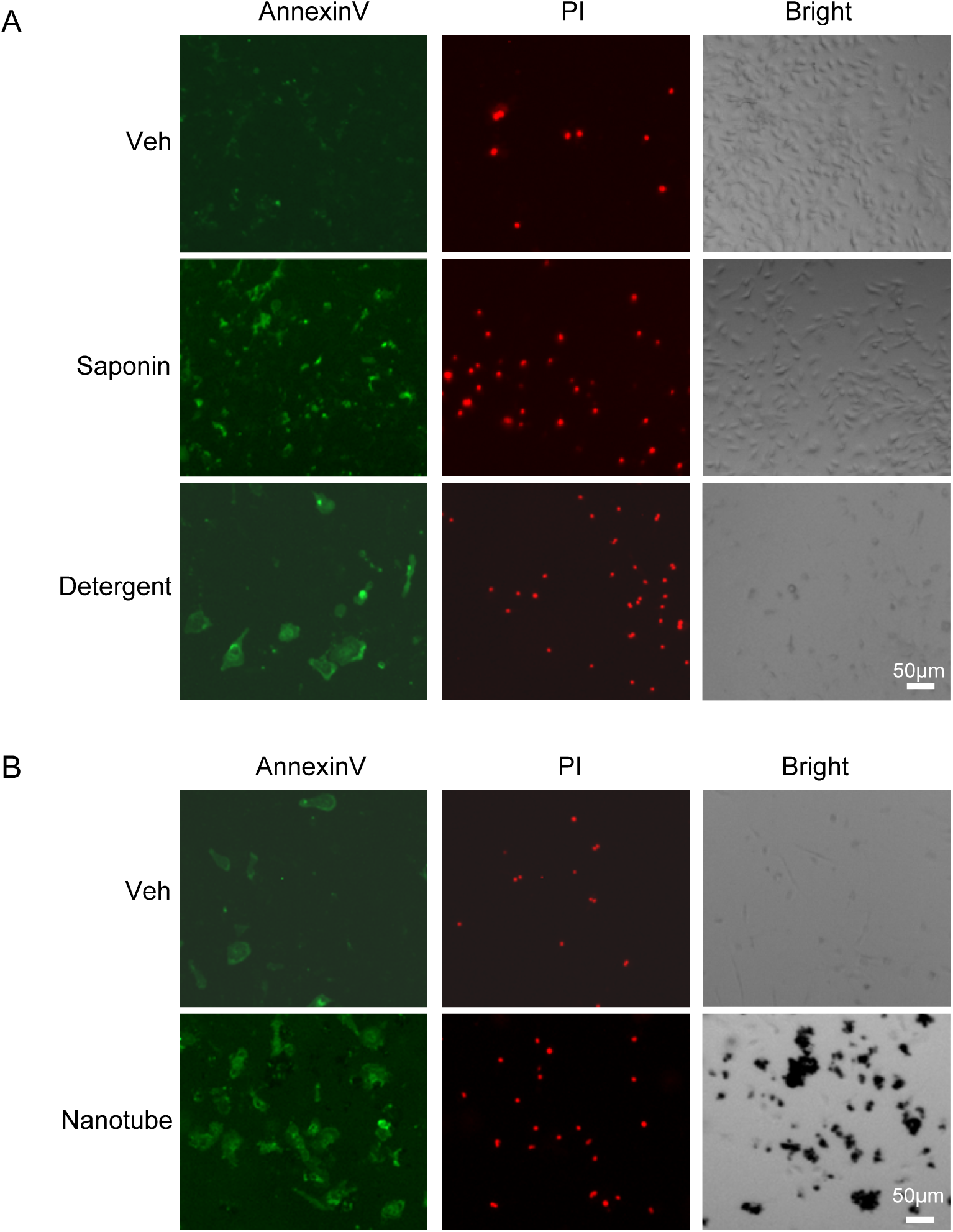
Environmental materials damage macrophage plasma membranes. (A) BMDMs were treated with saponin, detergent, or a vehicle. (B) BMDMs were treated with carbon nanotubes or a vehicle. Plasma membrane damage was assessed by Annexin V staining, and cell death was indicated by PI staining. Horizontal scale bar, 50 μm.

### Membrane damage by environmental materials promotes macrophage fusion

MGCs, a subset of functionally distinct multinucleate giant cells fused from macrophages, play critical roles in tissue homeostasis, host defense, and inflammatory responses. In situations such as granulomatous diseases or implant site reactions, MGCs contribute to the chronic inflammatory responses. Macrophage fusion is governed by various signaling pathways, including cytokine-driven mechanisms ^17–19,25,26^. Because the membrane damage is induced by environmental factors but not infections, we used a well-described condition that induces the formation of FBGCs by cytokines M-CSF and IL-4 ^27^. Early studies showed that needle-shaped nanoparticles promote MGCs in the lung and tumor microenvironment ^28,29^. As expected, treatment of BMDMs with carbon nanotubes enhanced the formation of FBGCs relative to treatment with the vehicle when IL-4 and M-CSF were supplied to the culture **(Figure 2A)**. Interestingly, treatment with surfactants saponin and detergent, both damaging macrophage membranes as carbon nanotubes **(Figure 1)**, also promoted the formation of FBGCs **(Figure 2B-C)**. These data suggest that membrane damage enhances macrophage fusion, a property of macrophages to respond to foreign materials.

**Figure 2.**
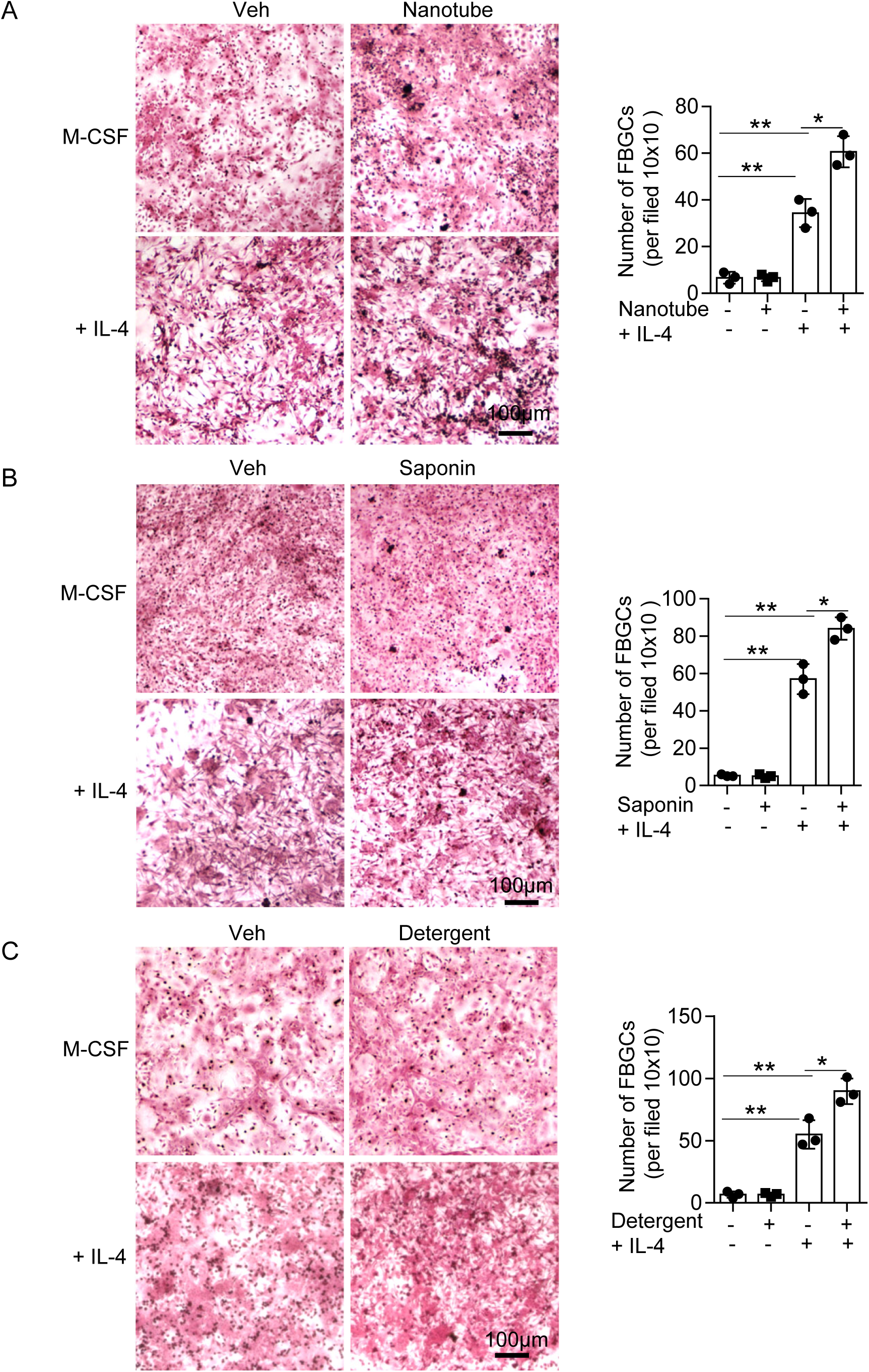
Environmental material-caused membrane damage promotes macrophage fusion. BMDMs were treated with carbon nanotubes (A), saponin (B), detergent (C), or a vehicle and then cultured with or without IL-4. Macrophage fusion was assessed by H&E staining. Horizontal scale bar, 100 μm. FBGC, foreign body giant cells; Veh, vehicle. Data (mean ± SD) shown are representative of 2independent experiments (n = 3 per group). Student’s *t*-test, *p < 0.05; **p < 0.005.

### Saponin-induced macrophage fusion is CD44-dependent

CD44, a cell surface adhesive molecule essential in cell-cell interactions, migration, and adhesion, plays a vital role in macrophage fusion ^19,26,30^. Because of the environmental materials interact with and damage the cellular membranes, we inferred that these materials interfere the function and even the expression of CD44 on the surface of macrophages. To test this, we treated macrophages with saponin and observed elevated levels of CD44 using flow cytometry followed by surface stain of CD44 **(Figure 3A)**. Next, to explore the role of CD44 in macrophage fusion induced by environmental damage, we generated *Cd44* knockout (KO) in BMDMs using a CRISPR-Cas9 system **(Supplemental Figure 2)** and treated the cells with or without IL-4 in the presence of M-CSF. As shown in Figure 2, without addition of IL-4, we did not observe cell-cell fusion in either *Cd44* WT or KO BMDMs in the presence or absence of saponin **(Supplemental Figure 3)**. Consistently, administration of IL-4 promoted MGC formation in WT BMDMs, whereas addition of saponin further enhanced this process, displaying greater numbers of MGCs, of which more cells contain 3 or more nuclei **(Figure 3B-C)**. Under this condition, the cell-cell fusion in *Cd44* KO BMDMs was significantly decreased, and the effects of saponin on macrophage fusion were diminished **(Figure 3B-C)**, suggesting that saponin-mediated enhancement of macrophage fusion is CD44-dependent.

**Figure 3.**
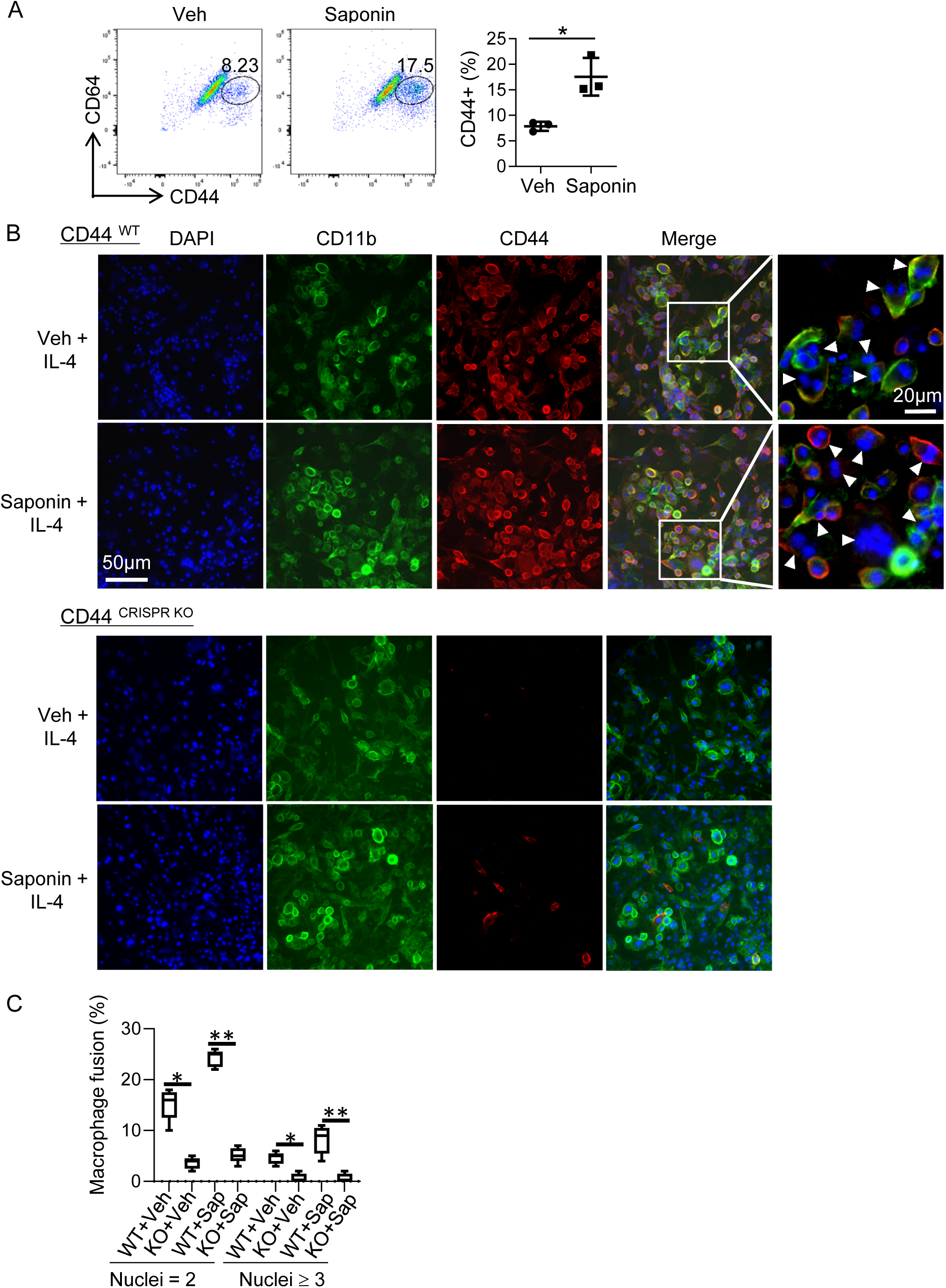
Saponin enhances macrophage fusion via upregulation of CD44. (A) Flow cytometry of CD44 expression on BMDMs after treatment with or without saponin. (B) Representative fluorescent images of *Cd44*-WT (Upper) or *Cd44*-deficient (Lower) BMDMs treated with saponin or a vehicle in the presence of IL-4. Images were acquired by using Cellomics. MGCs are indicated by arrowheads. (C) Statistical analysis of macrophage fusion in (B), respectively. One hundred cells (n = 100) were assessed in each sample. Percentages of cells with 2 or ≥3 nuclei were depicted. Veh, vehicle; Sap, saponin. Data shown are representative of 2 independent experiments. Student’s *t*-test, *p < 0.05.

### Saponin and detergent inhibit the expression of ROS in macrophages

ROS, one of the most important antimicrobial mediators produced by macrophages, serves as a major defense mechanism against pathogens. To understand the consequence of membrane damage caused by environmental materials on the defensive function of macrophages, we investigated the production of ROS in macrophages in response to these materials. BMDMs were treated with saponin, detergent or a vehicle and the levels of ROS was assessed by 2’,7’-dichlorodihydro-fluorescein diacetate (DCFH-DA) staining. Both saponin and detergent significantly decreased ROS levels compared to the vehicle (**Figure 4A-B, D-E**). As revealed by reverse transcription-polymerase chain reaction (RT-qPCR), treatment with saponin or detergent suppressed mRNA expression of macrophage-associated markers, such as pro-inflammatory cytokines (IL-1β, IL-6, and TNFα) and iNOS (encoded by *Nos2*) relative to the treatment with the vehicle (**Figure 4C-F**). These results indicate that exposure to environmental materials inhibits the expression of inflammatory mediators of macrophages, which may be the negative consequence of damage to plasma membranes because the membranes organize receptors, adaptor proteins, and kinases important for signal transduction.

**Figure 4.**
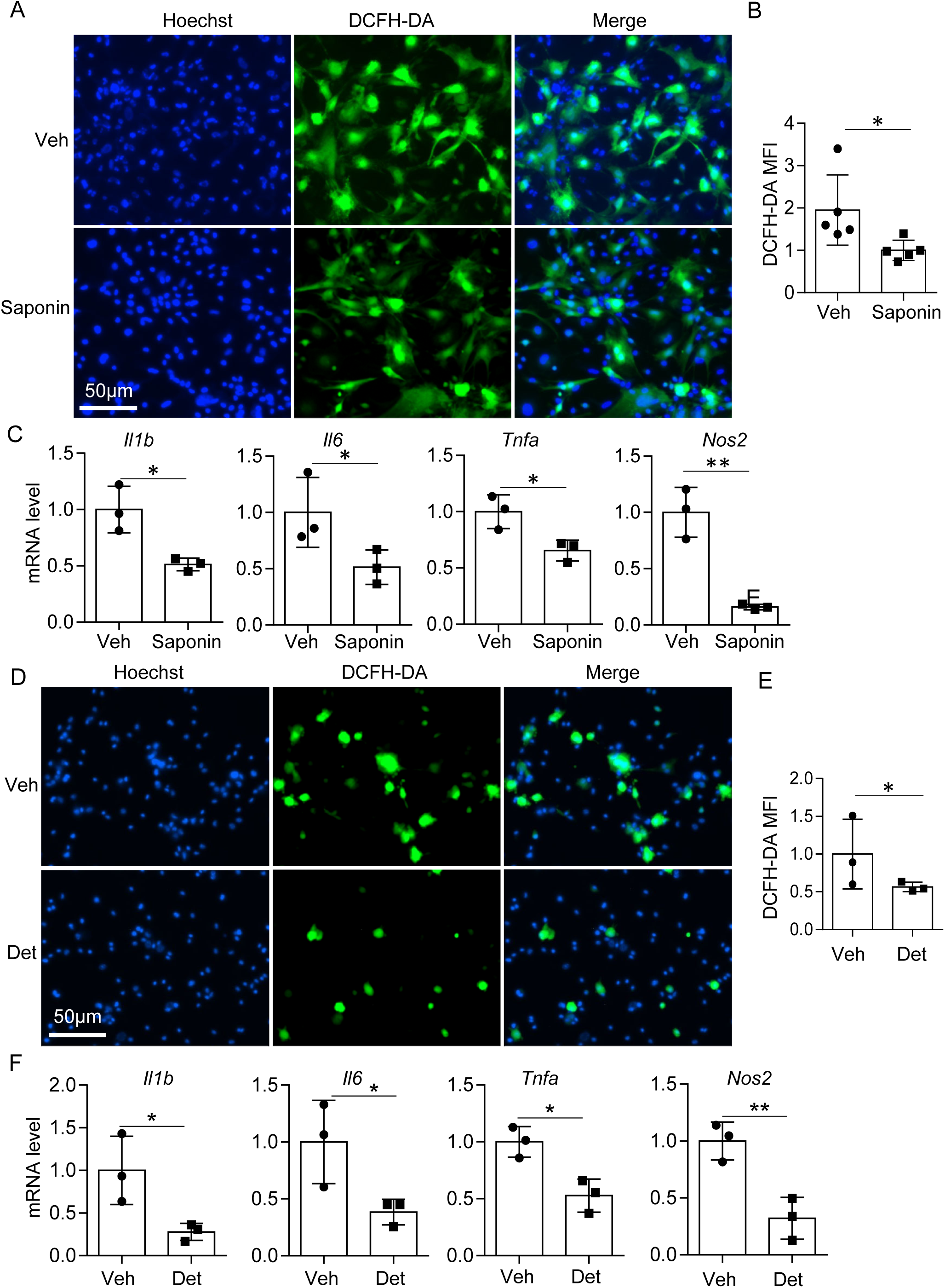
Environmental materials (saponin and detergent) decrease the production of ROS and pro-inflammatory cytokines in macrophages. BMDMs were treated with saponin (A-C), detergent (D-F), or a vehicle and stained with DCFH-DA and Hoechst. Veh, vehicle; Det, detergent. (A, D) Representative fluorescent images. Horizontal scale bar, 50 μm. (B, E) Mean fluorescent intensity (MFI) of DCFH-DA. (C, F) RT-qPCR of mRNA expression of indicated genes. *Actb* was used as an internal loading control. Data (mean ± SD) shown are representative of 2 independent experiments (n = 3 per group). Student’s *t*-test, *p < 0.05; **p < 0.005.

### Damage to macrophage membranes decreases macrophage phagocytosis

Macrophages are key players in the immune system through detecting and clearing pathogens through phagocytosis. This process is vital for host defense against a wide range of pathogens, including fungi and bacteria. To further understand how membrane damage influences the function of macrophages, we treated BMDMs with saponin for 10 minutes, and after washing, incubated the resulting macrophages with CFSE-labeled live *Escherichia coli* for 30 minutes. Following fixation, the cells were stained with DAPI to visualize macrophage nuclei. We found that treatment with saponin significantly decreased phagocytosis in macrophages compared to the vehicle-treated cells **(Figure 5A-B)**. We also assessed the impact of laundry detergent on macrophage phagocytosis. BMDMs were treated with detergent for 10 minutes, and after being washed, co-cultured with CFSE-labeled live *Candida albicans* for 30 minutes. Similar to phagocytosis of *E. coli*, treatment with detergent impaired phagocytosis of *C. albicans* in macrophages compared to the vehicle-treated cells **(Figure 5C-D)**. These results indicate that both detergent and saponin impair macrophage phagocytosis, which may be due to the membrane damage caused by the environmental factors. Our findings suggest that environmental exposures may impair immune functions and thus increase susceptibility to infections.

**Figure 5.**
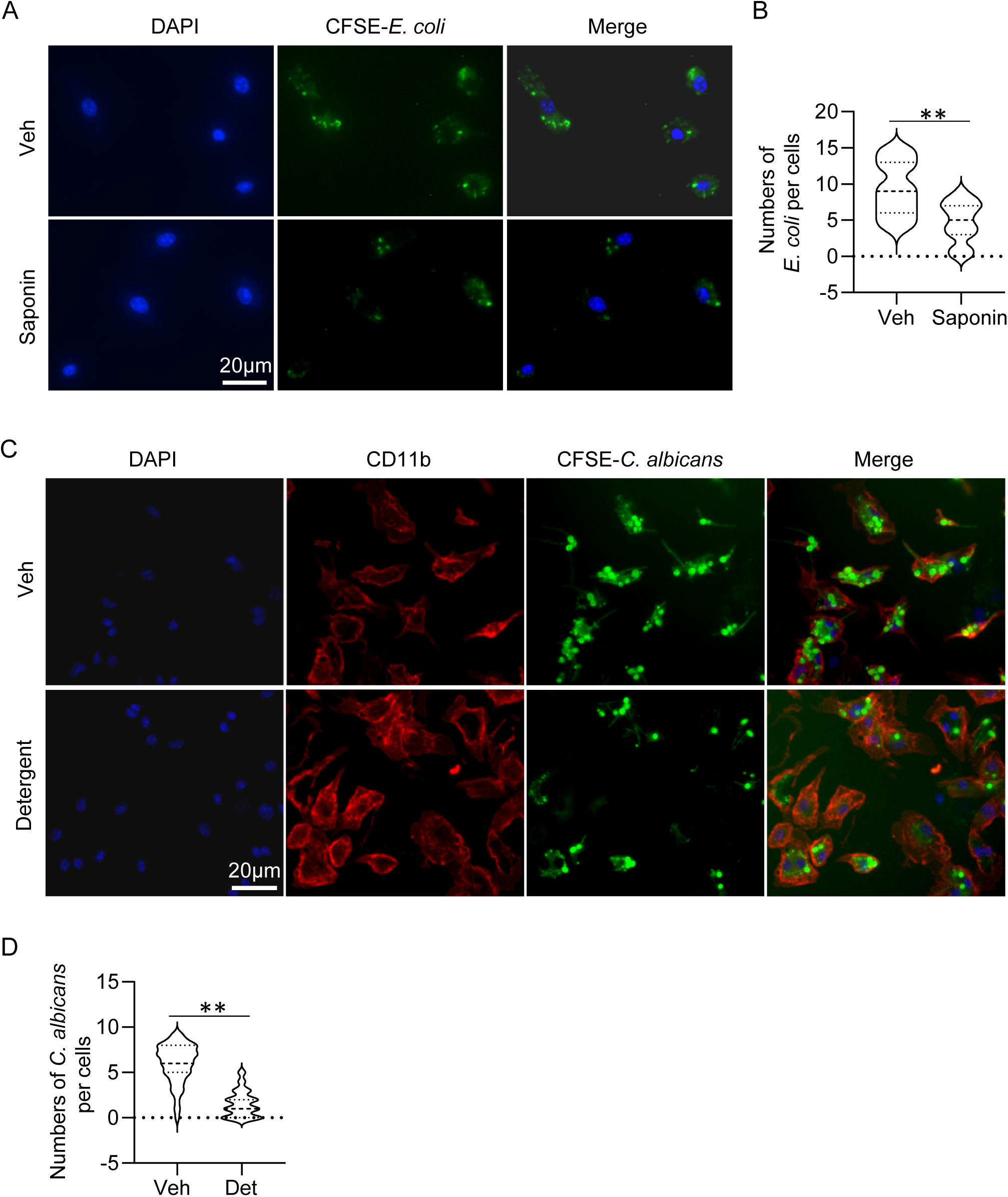
Environmental material-caused membrane damage impairs the phagocytic function of macrophages. BMDMs were cocultured with CFSE-labelled *E. coli* (A-C) or *C. albicans* (D, E). After fixation, the cells were stained with CD11b (red) and DAPI (blue). (A, C) Phagocytosis of *E. coli* (A) or *C. albicans* (C) by BMDMs treated with/without detergent or saponin, respectively. Horizontal scale bar, 20 μm. (C, D) Statistical analysis of (A, C). One hundred cells (n = 100) were assessed in each sample. Det, detergent. Data shown are representative of 3 independent experiments. Student’s *t*-test, **p < 0.005.

### Airway exposure to laundry detergent increases susceptibility to Candida infection in mice

To understand the effects of environmental hazards on host defense, we chose to examine laundry detergents, which are commonly used for cleaning clothes and linens and known to cause respiratory complications, including asthma ^9–12^, in a mouse model of Candida airway infection. Mice were intratracheally (i.t.) administered a laundry detergent or PBS, followed by a Candida i.t. challenge on the same day, once daily for four consecutive days. The mice were analyzed the next day after the last dose of Candida **(Figure 6A)**. Using colony-forming unit (CFU) assays, we found that the detergent-treated group had higher fungal burdens and lower body weights than the control group **(Figure 6B-C)**. Administration of detergent decreased the numbers of macrophages, eosinophils, and neutrophils in the BALF **(Figure 6D)**, suggesting that exposure to detergent impaired innate immune responses in the airway.

**Figure 6.**
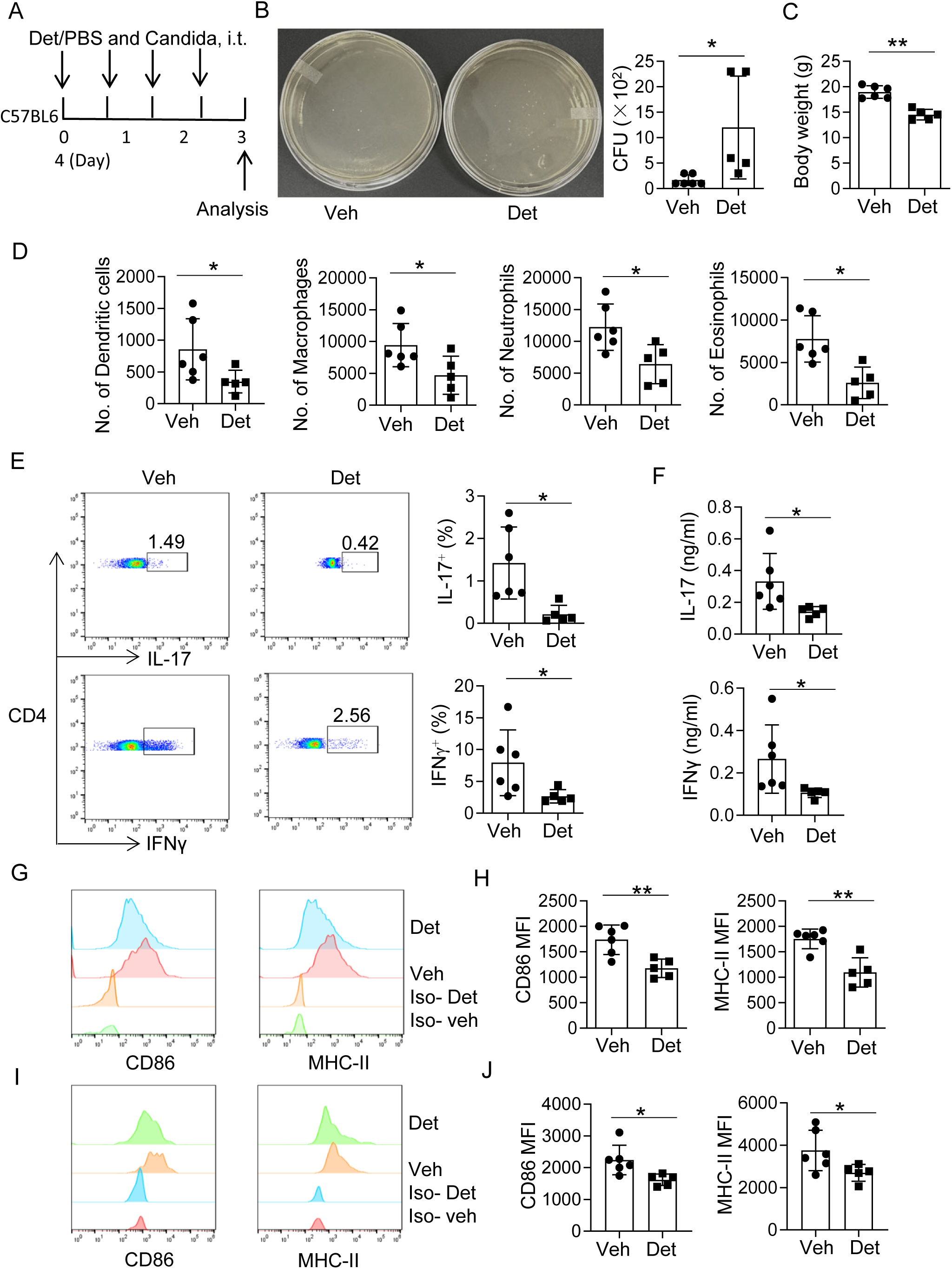
Laundry detergent increases susceptibility to Candida airway infection in mice. (A) C57BL/6 mice were intratracheally (i.t.) administered live Candida following i.t. receiving a laundry detergent or PBS once daily for four consecutive days, and sacrificed on the next day after the last dose of Candida. (B) Colony-forming unit (CFU) assays of Candida burden in the bronchoalveolar lavage fluids (BALF). (C) Body weights of the two groups of mice. (D) Profiles of myeloid cells in BALF. (E) Flow cytometry of IL-17- and IFNγ-expressing CD4^+^ TH cells in BALF. (F) ELISA of the levels of IL-17 and IFNγ in BALF. (G) Flow cytometry of MHC-II and CD68 on macrophages on a CD11b^+^Cd11c^lo^ gate. (H) Mean fluorescence intensity of MHC-II and CD68 in (G). (I) Flow cytometry of MHC-II and CD68 on dendritic cells on a CD11b^+^Cd11c^hi^ gate. (J) Mean fluorescence intensity of MHC-II and CD68 in (I). Veh, vehicle; Det, detergent; Iso, isotype control. Data (mean ± SD) are representative of two experiments. n = 5-6 per group. Student’s *t*-test, \**p*< 0.05; **p < 0.005.

Besides the innate immune profiles, exposure to detergent also impeded adaptive immune responses, including decreases in the frequencies of TH17 (IL-17^+^) and TH1 (IFNγ^+^) cells in the CD4^+^ fraction of BALF cells, and the levels of TH17 cytokine IL-17 and TH1 cytokine IFNγ in BALF **(Figure 6E-F)**. To understand how airway exposure to detergent influences T cell responses, we assessed the expression of CD86 and MHC-II by macrophages and dendritic cells, which are primarily involved in costimulation and antigen presentation for T cell activation and expansion. Interestingly, treatment with detergent not only damaged the membranes of macrophages (**Figure 1B**) but also inhibited the expression of CD86 and MHC-II by macrophages **(Figure 6 G-H)**. Similarly, exposure to detergent suppressed the expression of CD86 and MHC-II by dendritic cells (DCs) **(Figure 6 G-H)**. Collectively, treatment with laundry detergent suppresses the antigen presentation and costimulatory function of macrophages and DCs, leading to decreased immune responses in both innate and adaptive arms of the immune system, subsequently hindering host defense.

## Discussion

Macrophages, a critical compartment of the innate immune system, also bridge innate and adaptive immunity through recognition, phagocytosis, and digestion of pathogens and foreign materials, and presentation of antigens to T cells, mediating T cell activation and expansion. Environmental materials, such as tungsten carbide, nanotubes, and detergents, have been shown to influence macrophage function ^14–16^. However, how these hazards modulate macrophage function is not fully understood.

Cell membranes serve as a barrier separating the cell’s inner contents from its external environment and are responsible for cell-cell and cell-environment communications. Intensive studies have shown that various mechanisms, including interaction with environmental materials, excessive immune responses, and cell ageing, can cause cell membrane damage, leading to cell function alterations and even cell death ^31–33^. Because macrophages reside in the alveolar and intraepithelial spaces of the lung as well as the skin and gut, where they have the opportunity to interact with the environment, we hypothesize that the environmental materials cause damage to the membranes and subsequently impair the function of these cells. Indeed, our study has found that nanoparticles (WC, carbon nanotubes, and kaolin) and soluble surfactants (saponin and laundry detergent) can induce cell membrane damage in macrophages, but during the tested period, only slightly increase cell death **(Figure 1)**. Therefore, many environmental factors can disrupt the integrity of the plasma membrane of macrophages, and thus may lead to negative consequences in the functionality of these cells.

MGCs, formed from macrophage fusion, have unique functions in the immune system. In tuberculosis, MGCs form granulomas that constrain Mycobacterium tuberculosis when the bacteria escape lysosomes by preventing phagosome-lysosome fusion and can survive within the host cells ^34^. This is considered a defense mechanism to mitigate host tissue damage. Similarly, in the lungs of hard metal disease ^35–39^ and silicosis ^40^, macrophages fuse into MGCs; however, these MGCs fail to clear the ingested hard metal or silica particles, and the accumulation of MGCs results in deadly inflammation and tissue destruction. Whether cell membrane damage contributes to the pathogenesis of MGC diseases is unclear. Since there is a lack of infectious clues in exposure to environmental materials compared to pathogens, we examined the influence of macrophage membrane damage on the formation of FBGCs. Interestingly, both carbon nanotubes and surfactants (saponin and detergent) facilitate MGC formation **(Figure 2)**, although the underlying molecular mechanism remains unknown. CD44, a cell surface adhesive molecule, plays a vital role in macrophage fusion ^19,26,30^. We observed that saponin, one of the tested environmental hazards, upregulates the expression of CD44 on macrophages; KO of *Cd44* diminishes the effects of saponin on the stimulation of macrophage fusion **(Figures 3 and 4)**. In summary, environmental factors damage the membranes of macrophages and elevate the expression of CD44; these alterations together promote MGC formation.

As a defense mechanism, macrophages recognize the pathogen-associated molecular patterns (PAMPs) carried by pathogens through pattern recognition receptors (PRRs), such as Toll-like receptors (TLRs) and C-type lectin receptors (CLRs) that mediate the recognition of bacteria and fungi, leading to phagocytosis of these pathogens and formation of a phagosome. The phagosome containing engulfed pathogens then fuses with a lysosome. It forms a phagolysosome, where an array of active chemical materials [including reactive oxygen species (ROS) and nitric oxides] and enzymes digest and clear the phagocytosed pathogens. To determine whether environmental factors disrupt the defense mechanisms of macrophages, we evaluated the effect of surfactants on the expression of reactive oxygen species (ROS). Treatment with either saponin or detergent results in a decrease in the expression of ROS in macrophages **(Figure 4)**, suggesting that surfactants suppress lysosomal function in these cells, which is associated with membrane damage caused by the environmental materials. Furthermore, surfactants decrease the expression of mRNAs encoding pro-inflammatory cytokines (IL-1β, IL-6, and TNFα) and the inducible nitric oxide synthase (INOS or Nos2) **(Figure 4)**, weakening the defensive and inflammatory functions of macrophages. Although we cannot rule out that these surfactants may enter the cells and disrupt the cell function via interaction with intracellular signal components, the membrane damage caused by these materials contributes to the disturbance of the incoming signals, given many receptors, kinases, and signal adaptor proteins are anchored in or docked inside the plasma membranes. As discussed above, macrophages utilize PRRs to recognize pathogens. Because of this, we inferred that membrane disruption by the environmental materials impairs the phagocytic function of macrophages. Our research confirmed that treatment with both saponin and detergent dramatically decreases the phagocytosis of *E. coli* and *C. albicans*, respectively, by macrophages **(Figure 5)**, which is associated with cellular membrane damage of these cells **(Figure 1)**.

Exposure to various environmental materials through the airway can lead to lung complications. Inhalation of PM, such as carbon nanotubes, can cause airway inflammation and lung damage ^7,8^, whereas exposure to detergent residues can result in airway constriction and hyperresponsiveness, increasing the risk of asthma skin ^9,10^. We also found that surfactants inhibit the expression of ROS and decrease the phagocytic function of macrophages (see discussion above). However, it is not clear whether and to what extent these environmental toxins influence host defense. To further evaluate this, we used a Candida airway infection mouse model. Airway exposure to detergent increases fungal burdens in the lung, associated with decreased airway infiltration of myeloid cells, including dendritic cells, macrophages, neutrophils, and eosinophils **(Figure 6B, D)**, of which macrophages and neutrophils play an essential role in anti-fungal immunity ^41^. Moreover, treatment with detergent also suppresses adaptive immunity, manifesting decreased frequencies of TH1 and TH17 cells and their signature cytokines, IFNγ and IL-17, respectively, in the BALF **(Figure 6E-F)**. T cell responses rely on antigen-presenting cells, including DCs and macrophages, to present antigens through their MHC complexes (for TH cells, MHC-II) and to provide costimulation through the B7 molecules. Airway exposure to detergent inhibits the expression of CD86, a member of the B7 family, and MHC-II on macrophages and DCs **(Figure 6G-J)**. In line with the decreases in the expression of pro-inflammatory cytokines (IL-1β, IL-6, and TNFα) in macrophages **(Figure 4)**, inhibition of CD86 and MHC-II leads to a defect in TH1 and TH17 cell responses. Taken together, airway exposure to detergent results in the decline of both innate and adaptive arms of the immune system, therefore impairing host defense to fungal airway infection.

In summary, our data suggest that environmental materials cause a broad spectrum of negative effects on the immune system, increasing susceptibility to infections. The results of this study may offer novel insights into the development of preventative and therapeutic options.

## Materials and Methods

### Mice

All mice were housed in the specific pathogen-free animal facility, and all procedures were conducted in accordance with protocols approved by the Institutional Animal Care and Use Committee of the University of New Mexico Health Sciences Center.

### Administration of laundry detergent and *C. albicans* airway infection

Sex-matched C57BL/6 mice (6–8 weeks old) were intratracheally (i.t.) administered 40 μL of a commercially available liquid laundry detergent (1%) or vehicle control (phosphate-buffered saline, PBS). Two hours later, the mice were i.t. challenged with 1 × 10^6^ *C. albicans in* 40 μL of PBS, daily for four consecutive days. Mice were sacrificed one day after the final dose of *C. albicans*, and lung tissues and bronchoalveolar lavage fluid (BALF) were collected for further analysis. No significant differences were observed between male and female mice.

### Colony formation assay

One percent (10 μl out of 1 ml) of each BALF sample was plated onto a yeast extract peptone dextrose (YPD) agar plate and incubated at 30°C overnight. Colonies were counted based on size and morphology, with large white colonies quantified as *C. albicans*.

### Generation and culture of BMDMs

C57BL/6 mice (6–8 weeks old) were euthanized by cervical dislocation. Femurs and tibias were harvested, and bone marrow (BM) cells were flushed from the medullary cavities. BM cells were cultured in RPMI-1640 medium (Corning) supplemented with 2 mM L-glutamine (Gibco), 10% fetal bovine serum (FBS; Atlanta Biologicals), and 100 U/ml penicillin-streptomycin (Gibco) in the presence of 20 ng/ml recombinant mouse macrophage colony-stimulating factor (M-CSF; Biolegend). Cultures were incubated at 37°C with 5% CO2 for 6–7 days to induce macrophage differentiation. BMDMs were harvested using Accutase® Cell Detachment Solution (Biolegend). To form MGCs, BMDMs were supplied with 10 ng/ml IL-4 (PeproTech) in the presence of M-CSF (20 ng/ml) and cultures as above for 4 days ^27^.

### Treatments with environmental materials and assessment of plasma membrane damage

BMDMs were treated with 0.1% saponin (S7900, Sigma-Aldrich), 1% liquid laundry detergent (a common commercial brand), or a vehicle for 10 min, or 10 μg/ml carbon nanotube (10-30 nm, #900-1201, SES Research), 10 μg/ml kaolin (K7375, Sigma-Aldrich), WC (150-200 nm, 778346, Sigma-Aldrich), or a vehicle overnight. The resulting cells were labeled with Alexa Fluor 488-conjugated Annexin V to identify plasma membrane disruption and propidium iodide (PI) to identify dead cells. Images were acquired by using the EVOS FL Cell Imaging System (Thermo Fisher Scientific).

### Hematoxylin and eosin (H&E) staining

BMDMs were treated with saponin (12.5 μg/ml), carbon nanotubes (1.25 μg/ml), laundry detergent (1%), or PBS (vehicle) in the presence of M-CSF or IL-4 for 6 days. After culture, the cells were fixed with 4% paraformaldehyde at room temperature for 10 min and stained with H&E according to standard protocols. Images were acquired by using the EVOS FL Cell Imaging System (Thermo Fisher Scientific).

### Flow Cytometry

Antibodies against CD45.2 (104), CD4 (RM4.5), CD11b (M1/70), CD11c (N418), Ly-6G (1AB-Ly6g), SiglecF (E50-2440), Gr-1 (RB6-8C5), IFNγ (XMG1.2), IL-17A (eBio17B7), CD86 (GL-1), MHC-II (M5/114.15.2) were purchased from eBioscience. Cells stained with indicated markers were analyzed on an Attune NxT Flow Cytometer (Thermo Fisher Scientific), and the data were processed by Flowjo software (FlowJo, LLC).

### Culture and CFSE-labelling of *C. Albicans* and *E. coli*

*C. albicans* SC5314 (ATCC, MYA-2876), recovered from a −80°C glycerol stock, was grown in YPD medium (L22052801, US Biological Life Sciences) for 16 h at 30°C in an orbital shaker at 250 rpm to mid-log phase. *E. coli* was cultured in LB medium for 16 h at 37°C with orbital shaking at 250 rpm to mid-log phase. After being washed, *C. albicans* or *E. coli* were suspended and incubated in 10 µM CFSE (C34554, Invitrogen)-PBS in the dark at room temperature for 10 min. The cells were then washed with cold PBS, suspended in RPMI medium, and stored on ice for later use.

### Phagocytosis assay

BMDMs, pre-treated with detergent, saponin, or vehicle as above, were incubated with CFSE-labelled live *C. albicans* or *E. coli* [Macrophage to *C. albicans* (or *E. coli*) = 1:10] at 37°C in an atmosphere containing 5% CO2. After 30-min co-culture, the plate was washed with cold PBS. The resulting BMDMs were fixed with 4% paraformaldehyde for 15 min and stained with DAPI and CD11b (M1/70) for 10 min. High-content microscopy with automated image acquisition and quantification was carried out using a Cellomics HCS scanner and iDEV software (Thermo Fisher Scientific). All data collection, object assignments, identification of regions of interest, and analyses were performed by the software in an unsupervised manner.

### DCFH-DA staining

2 × 10^5^ BMDMs were plated in a 24-well plate and cultured overnight. The next day, the cells were treated with 0.1% saponin, 1% liquid laundry detergent, or a vehicle for 10 min. After being washed with cold PBS, the resulting cells were stained with DCFH-DA (Cayman Chemical) and Hoechst at 37 °C for 30 min. Following washes, the cells were kept in PBS, and images were acquired using a Cellomics HCS scanner equipped with iDEV software (Thermo Fisher Scientific). The cells were then lysed in 200 μl of radioimmunoprecipitation assay (RIPA) buffer, and 100 μl of each lysate was transferred into a black 96-well plate for measuring fluorescence intensity using a fluorescence microplate reader at an excitation wavelength of 485 nm and an emission wavelength of 530 nm. Protein concentrations were detected in 1 μl of lysates diluted in 100 μl of 1× protein assay solution using the Quick Start™ Bradford 1x Dye Reagent (Bio-Rad). Fluorescence intensities were normalized to the protein concentrations as described ^42^.

### Generation of CRISPR knockout (KO)

CRISPR KO of CD44 in macrophages was performed as described ^43^. Briefly, gRNAs (synthesized by Integrated DNA Technologies) were mixed with Cas9 (IDT Alt-R S.p. Cas9 Nuclease V3) at a molar ratio of 2:1 and incubated at room temperature for 20 min. 5 x 10^6^*in-vitro* differentiated BMDMs were washed and resuspended in 20 μl of P3 primary nucleofection solution (V4XP-3024, Lonza). The cells were then added to the Cas9-gRNAs complex and mixed by pipetting. The mixture was nucleofected using the CM-137 condition on a 4D-Nucleofector (4D-Nucleofector Core Unit, AAF-1002B; 4D-Nucleofector X Unit, AAF-1002X, Lonza). The resulting cells were maintained in the presence of M-CSF, and KO efficiency was assessed by RT-qPCR.

### RT-quantitative (q) PCR

Gene mRNA expression was determined by qPCR using the CFX Connect Real-Time PCR Detection System (Bio-Rad Laboratories) following reverse transcription with SuperScript Reverse Transcriptase (18064022, Thermo Fisher Scientific). The primers were, *Nos2* forward, 5’-GGCAGCCTGTGAGACCTTTG, reverse, 5’-CGTTTCGGGATCTGAATGTGA ^44^; *Actb* forward, 5’-GACGGCCAGGTCATCACTATTG, reverse, 5’-AGGAAGGCTGGAAAAGAGACC ^45^; *Il6* forward, 5’-GAGGATACCACTCCCAACAGAC, reverse, 5’-AGCTATGGTACTCCAGAAGACC; *Il1b* forward, 5’-AACCTTTGACCTGGGCTGTC, reverse, 5’-AATGGGAACGTCACACACCA, and *Tnfa* forward, 5’-AGGCACTCCCCCAAAAGATG, reverse, 5’-TTTGCTACGACGTGGGCTAC. Data were normalized to a reference gene *Actb*.

### Statistical analysis

Data are expressed as mean ± SD. Statistical significance between groups was determined using an unpaired Student’s *t*-test. *P* ≤ 0.05 was considered statistically significant.

## Data availability

All data generated or analyzed during this study are included in this article.

## Acknowledgments

This work was supported in part by NIH grants HL148337 and AI187912 (X.O.Y.), DK132643 (M.L.), ES026673 (M.C.), and ES035780 (X.X.). We acknowledge the University of New Mexico Comprehensive Cancer Center Flow Cytometry Facility and Health Sciences Center Autophagy, Inflammation and Metabolism in Disease Center Core Facility (supported by NIH CA118100 and GM121176, respectively).

---

**Supplemental Figure 1.**
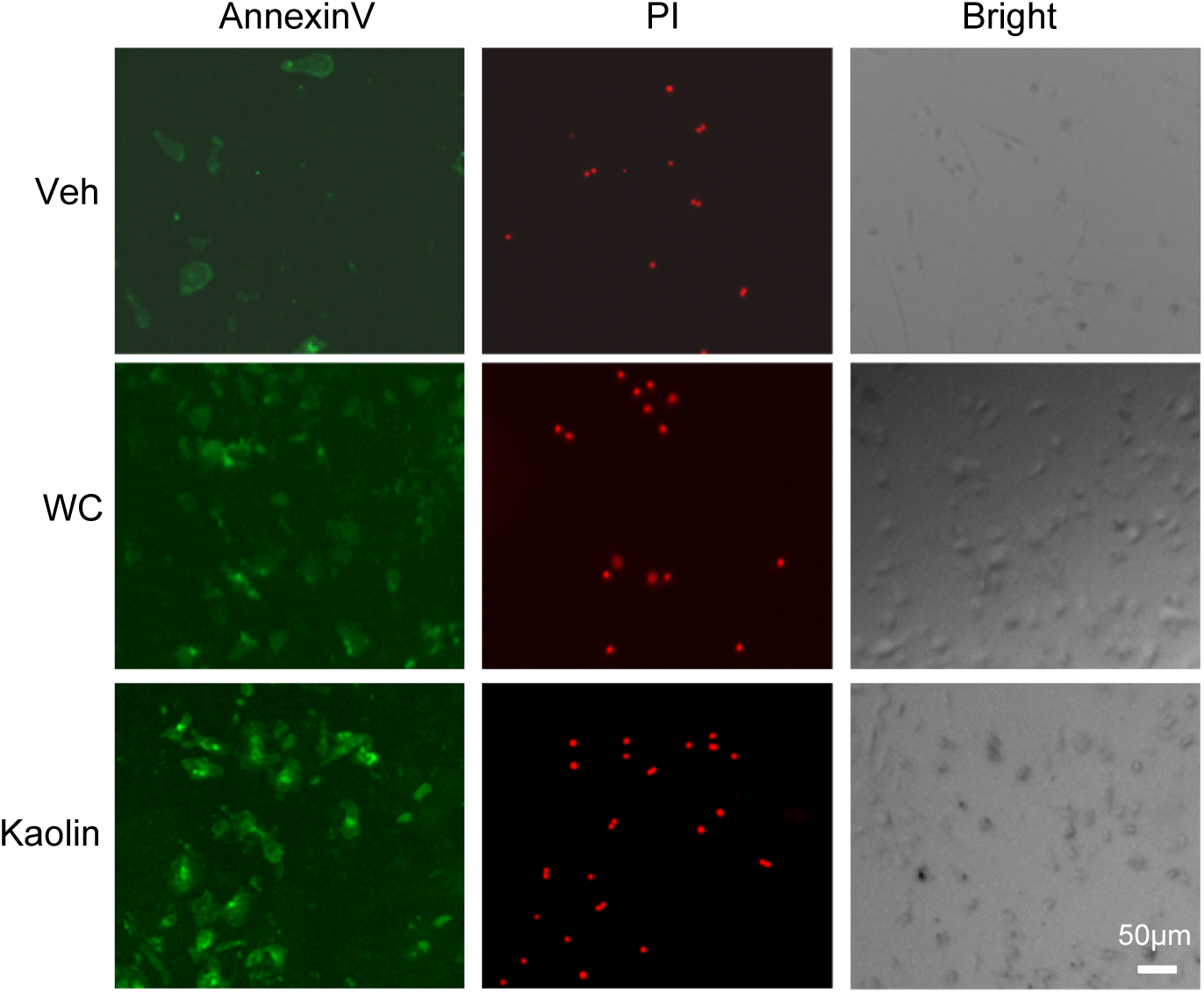
Tungsten carbide (WC) and kaolin damage macrophage plasma membranes. BMDMs were treated with WC, kaolin, or a vehicle (Veh). Plasma membrane damage was assessed by Annexin V staining, and cell death was indicated by PI staining. Horizontal scale bar, 50 μm. Note: The images of the vehicle control are the same as Figure 1B.

**Supplemental Figure 2.**
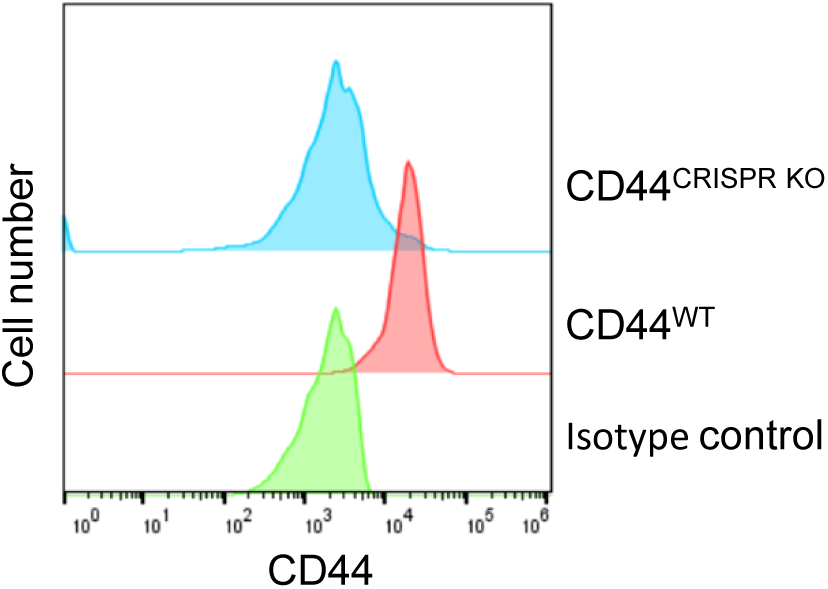
CRISPR-Cas9 knockout of *Cd44*. Flow cytometry of CD44 expression on BMDMs with or without CRISPR-Cas9 knockout of *Cd44*.

**Supplemental Figure 3.**
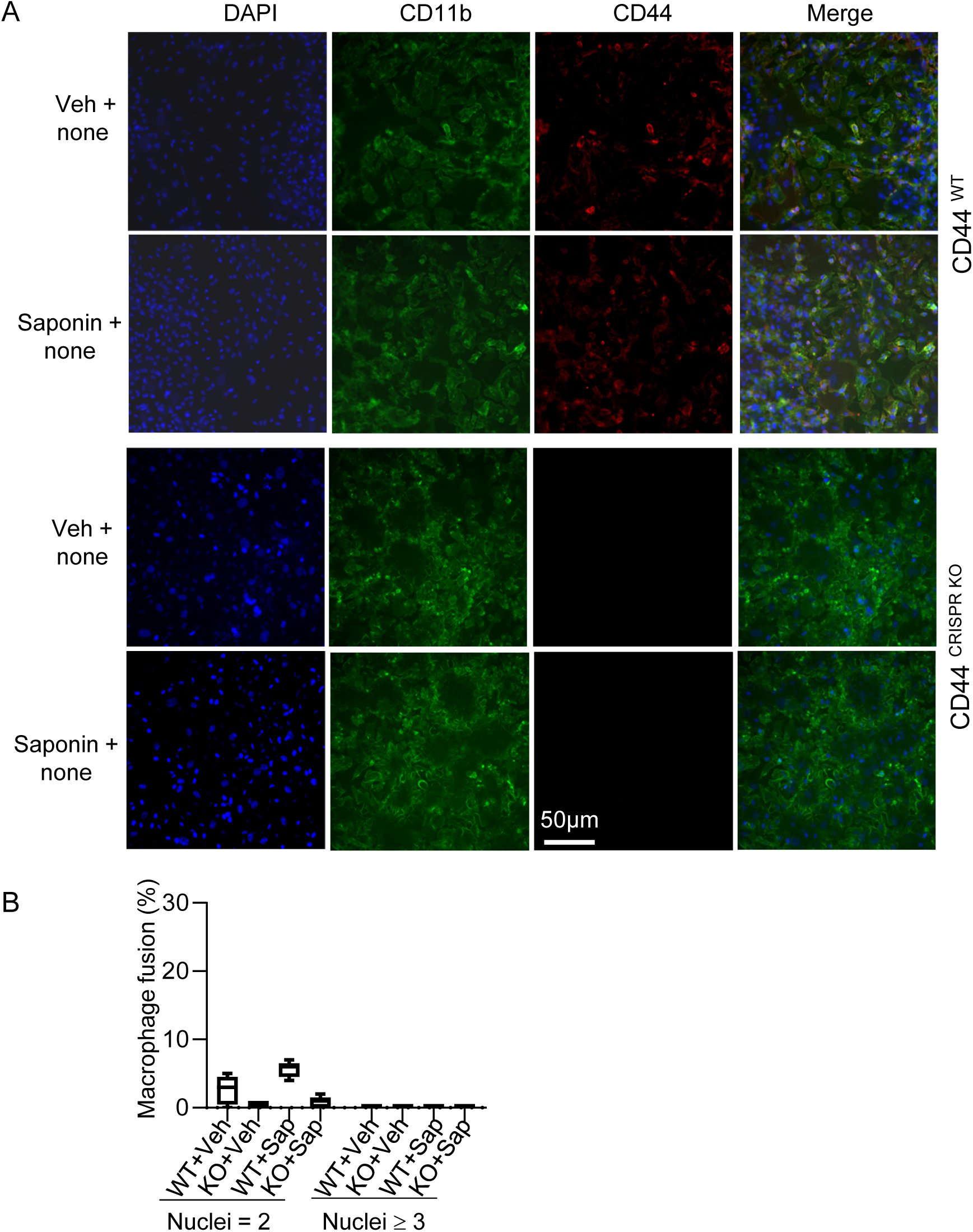
Effects of saponin on macrophage fusion in the absence of IL-4. (A) Representative fluorescent images of *Cd44*-WT (Upper) or *Cd44*-deficient (Lower) BMDMs treated with saponin or a vehicle in the absence of IL-4. Images were acquired by using Cellomics. (B) Statistical analysis of macrophage fusion in (A), respectively. One hundred cells (n = 100) were assessed in each sample. Percentages of cells with 2 or ≥3 nuclei were depicted. Veh, vehicle; Sap, saponin. Data shown are representative of 2 independent experiments.

